# Long-term Motor Learning in the Wild with High Volume Video Game Data

**DOI:** 10.1101/2021.09.15.460516

**Authors:** Jennifer B. Listman, Jonathan S. Tsay, Hyosub E. Kim, Wayne E. Mackey, David J. Heeger

**Affiliations:** Statespace Labs, Inc., New York, NY, USA; Department of Psychology, University of California, Berkeley, Berkeley, CA, USA; Helen Wills Neuroscience Institute, University of California, Berkeley, Berkeley, CA, USA; Department of Physical Therapy, University of Delaware, Newark, DE, USA

**Keywords:** esports, motor learning, video games, sensorimotor performance, motor acuity

## Abstract

Motor learning occurs over long periods of practice during which motor acuity – the ability to execute actions more accurately, precisely, and within a shorter amount of time – improves. Laboratory-based motor learning studies are typically limited to a small number of participants and a time frame of minutes to several hours per participant. Thus, there is a need to assess the generalizability of theories and findings from lab-based motor learning studies on much larger samples across longer time scales. In addition, laboratory-based studies of motor learning use relatively simple motor tasks which participants are unlikely to be intrinsically motivated to learn, limiting the interpretation of their findings in more ecologically valid settings. We studied the acquisition and longitudinal refinement of a complex sensorimotor skill embodied in a first-person shooter video game scenario, with a large sample size (N = 7174 participants, 682,564 repeats of the 60 sec game) over a period of months. Participants voluntarily practiced the gaming scenario for as much as several hours per day up to 100 days. We found improvement in performance accuracy (quantified as hit rate) was modest over time but motor acuity (quantified as hits per second) improved considerably, with 40-60% retention from one day to the next. We observed steady improvements in motor acuity across multiple days of video game practice, unlike most motor learning tasks studied in the lab that hit a performance ceiling rather quickly. Learning rate was a nonlinear function of baseline performance level, amount of daily practice, and to a lesser extent, number of days between practice sessions. In addition, we found that the benefit of additional practice on any given day was non-monotonic; the greatest improvements in motor acuity were evident with about an hour of practice and 90% of the learning benefit was achieved by practicing 30 minutes per day. Taken together, these results provide a proof-of-concept in studying motor skill acquisition outside the confines of the traditional laboratory and provide new insights into how a complex motor skill is acquired in an ecologically valid setting and refined across much longer time scales than typically explored.

## Introduction

Motor learning, the process of acquiring and refining skilled movements, is an essential human capacity (Du et al., 2021; Fitts & Posner, 1979; Kim et al., 2020; Krakauer et al., 2019; Reza Shadmehr et al., 2010). For instance, motor learning enables babies to acquire walking and talking skills, and athletes to become champions. These motor skills are honed over a lifetime of practice (Crossman, 1959; Ericsson et al., 1993), often demanding tremendous grit and intrinsic motivation (A. Duckworth, 2016; A. L. Duckworth et al., 2007; Mathis, 1967; R. M. Ryan & Deci, 2000).

In 1968, Fitts and Posner proposed a highly influential conceptual model of how motor learning progresses through three distinct stages (Fitts & Posner, 1979): cognitive, associative, and autonomous. The cognitive stage entails learning what movements to perform. This stage is typically short yet effortful (Adams, 1982; Dickinson & Weiskrantz, 1985), and typically produces the steepest improvements in performance, as task-specific strategies are learned (Daw et al., 2005). The association stage is characterized by learning to fine-tune how these movements should be performed. This stage is less effortful but occurs over a long period. Learners eventually reach the autonomous stage, when movements are performed not only accurately and rapidly, but also automatically and effortlessly.

The association stage yields the greatest improvements in motor acuity – the ability to execute actions more accurately, more precisely, and within a shorter amount of time (McDougle & Taylor, 2019; Müller & Sternad, 2004; Shmuelof et al., 2012, 2014; Wilterson, 2021). For instance, a baseball player who has learned all the fine-grained mechanical details of a baseball swing still requires continuous practice to consistently generate enough power to hit home runs. Movement kinematics, like the trajectory of the swing and the pivoting of the trunk, also improve in the association stage, becoming less variable and more coordinated over time (Deutsch & Newell, 2004; Guo & Raymond, 2010; Hung et al., 2008; Liu et al., 2006; Logan, 1988; Müller & Sternad, 2009).

Despite evidence for its importance, there is a paucity of studies examining improvements in motor acuity during the association stage of motor learning, a gap likely linked to the tight resource constraints on laboratory-based studies. Specifically, since in-lab studies are typically on the order of minutes to hours, studies on improvements in motor acuity that occur on the order of days to weeks (and even years) are rare (but see: (Berlot et al., 2020; Hardwick et al., 2019; Huberdeau et al., 2019; Semrau et al., 2012; Stafford & Dewar, 2014; Stratton, 1896; Wilterson & Taylor, 2021)). Moreover, the handful of studies that examine motor acuity have used relatively simple motor tasks (Flatters et al., 2014), like drawing circles as fast as possible within a pre-defined boundary (Shmuelof et al., 2012), throwing darts (Martin et al., 1996), or center-out reaching and grasping (Jordan & Rumelhart, 1992; R. Shadmehr & Mussa-Ivaldi, 1994). Whether insights generated in the lab can generalize to more naturalistic, ecological motor skills remains to be seen.

How then can we study the refinement of ecological, complex motor skills over longer timescales? There have been recent efforts to study motor skill acquisition outside the lab environment (Bönstrup et al., 2020; Drazan et al., 2021; Haar et al., 2021; Johnson et al., 2021; Stafford & Dewar, 2014; Tsay, Ivry, et al., 2021). Online video game platforms, in particular, provide an unprecedented opportunity to reach more than 400 million intrinsically motivated people who voluntarily hone their cognitive and motor skills over weeks, months and even years (Eichenbaum et al., 2014; Green & Bavelier, 2003). Specifically, first person shooter games, where players use tools and weapons as intuitive extensions of themselves in a virtual 3D environment, provide an ideal testbed for studying motor skill acquisition over a long duration of practice (Stafford & Vaci, 2021). Successful shooters are those who can efficiently identify and localize relevant visual targets (Green & Bavelier, 2006), as well as rapidly and accurately hit their targets – a refined motor skill. Mining the wealth of motor performance metrics from these first-person shooter games, we may be able to better understand how complex motor skills are developed and refined over longer timescales.

We have developed a video game, Aim Lab, that trains and assesses players to optimize their visuomotor performance in first-person shooter gaming scenarios. We have amassed a sizable dataset (> 20 million players) with a wide range of abilities from beginner to professional esports athlete, from which we sampled for this study (N = 7174 participants with over 62,170 game days of data). Most critical to this study, we analyzed how motor acuity improved over one hundred days of gameplay, providing a proof-of-concept in studying motor skill acquisition outside the confines of the traditional laboratory.

## Materials and Methods

### Apparatus

Aim Lab was written in the C# programming language using the Unity game engine. Unity is a cross-platform video game engine used for developing digital games for computers, mobile devices, and gaming consoles (Brookes et al., 2020). Players download Aim Lab to their desktop or laptop PC. Players control their virtual weapon in Aim Lab tasks, using a mouse and keyboard, while viewing the game on a computer screen. Performance data are uploaded to Aim Lab secure servers.

Aim Lab includes over ninety different scenarios for skill assessment and training. Each exercise is tailored to a facet of first-person shooter gameplay, and can be customized to prioritize accuracy, speed, and other basic components of performance. Various scenarios assess and train visual detection, motor control, tracking moving targets, auditory spatial-localization, change detection, working memory capacity, cognitive control, divided attention, and decision making. Aim Lab players can perform a variety of tasks or play the same task repeatedly. It is common for players to log in more than once on a given day to play.

### Task

Of all task options, the “Gridshot” task has the highest median number of repeats per player within the same day. This sixty-second task assesses motor control of ballistic movements. Three targets are presented simultaneously, at any given time, with a new target appearing after each target is destroyed. All targets are the same size, ranging between 1.3 - 1.7° (deg of visual angle), depending on viewing distance and field of view set by the player. New target appearance locations are randomized to one of twenty-five positions in a 5 × 5 grid, ranging between 4.8 - 9.1° wide and 5.1 – 7 .8° high, again depending on viewing distance and field of view which can be set by the player. The player destroys a target by moving their mouse to aim and clicking the left mouse button to shoot (Figure 1). Because multiple targets are present at once, combined with unlimited target duration and no explicit incentive to destroy any specific target, the player themself must decide the order in which to destroy the targets.

**Figure 1.**
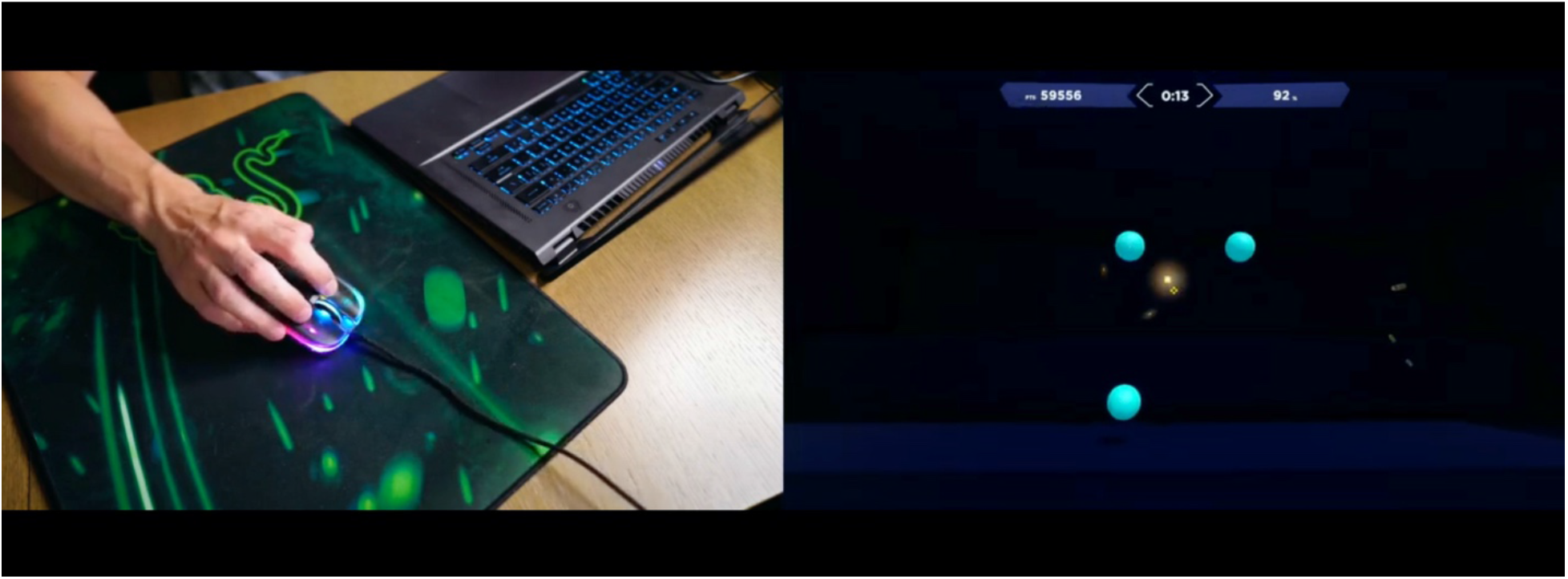
Gridshot task in Aim Lab. The 60-second run of Gridshot with three targets visible at all times is presented to a player on their computer screen (right) while the player controls on-screen movements and shots with a standard computer mouse (left).

Players receive immediate feedback upon target destruction; an explosion sound is emitted, and the orb-shaped target splinters into multiple pieces and then disappears. Players receive summary feedback after each sixty-second run of Gridshot, including score, hits per second (number of targets successfully destroyed per second), and hit rate (% of shot attempts that successfully hit a target). Points are added to the score when targets are hit and subtracted for shots that miss a target, and score is displayed at the top of the screen throughout the run. The number of points added for each target hit are scaled by time since the previous target hit. That is, more points are added when the time from the previous hit is shorter. Thus, players are incentivized to shoot targets rapidly and accurately as well as quickly plan their next movement and shot. While players are shown multiple metrics at the end of each run, it is likely that they are consciously optimizing for increased score. Some players compete for high scores on a leaderboard, which displays top scores across all players.

### Participants

Historical (pre-existing) longitudinal data were sampled from a subset of 100,000 randomly selected Aim Lab player IDs from those who signed up for an account on or after 7/1/20 and who played on more than one date between then and 1/30/21. Players are located throughout North, Central, and South America, Africa, Europe, Asia, Australia, and The Middle East. Aim Lab does not collect personal or demographic information from players. Based on an in-game survey completed by 4700 Aim Lab players (not necessarily overlapping with those whose data were analyzed here), we estimate that the player base is approximately 10% female and 90% male, with a median age of 18 years and range of 13-70 years.

Players were not required or requested to play any specific Aim Lab tasks; data were generated through typical use of the software on the players’ own gaming equipment in the setting and times of their choosing, likely at home. Play data were initially acquired for commercial purposes, are stored separately from player account data and without personal identifiers; thus, informed consent is not required for this study (Advarra Online ICF Pro00035991, Approved Apr 6, 2020).

### Data Analysis

We downloaded all Gridshot data for runs logged by the randomly selected players between 7/1/2020 – 1/30/21. Data for each sixty-second run of Gridshot included player ID (a unique code for each player), date and time, mode (practice or compete), score, hits per second, and hit rate.

We limited our analysis to runs played with the weapon type “pistol”, the most popular weapon used, and the analysis was limited to players who played more than one day. Game days greater than 100 and with fewer than three Gridshot runs were removed. We removed Gridshot runs with missing values or possible equipment problems (ex: date/time stamps in the future), cheating (proprietary Aim Lab methods for identifying cheaters), or unreasonably low scores (score ≤ 80% of the player’s median score on their first day playing Gridshot).

Our inclusion criteria resulted in 62,170 days of play from 7174 players totaling 682,564 Gridshot runs (60 sec each). Because score is a function of both hits per second and hit rate, we limited our analyses to hits per second and hit rate. For each player we calculated:

- *K* runs total (between 7/1/20 and 1/30/21)
- *M* days played (between 7/1/20 and 1/30/21)
- *K* runs per day
- Median of *K* runs per day, based on all days played

For each player’s first day playing Gridshot we calculated:

- Start date (calendar date of first run of Gridshot)
- Baseline score (from run 1 on game day 1)
- Baseline hit rate (from run 1 on game day 1)
- Baseline hits per second (from run 1 on game day 1)

On each successive game day we calculated:

- Days from last play (*M* days between calendar dates of play)
- Day number (the *m*^th^ day the player played the game)
- Score, hit rate, and hits per second for each run

Gridshot can be played in two modes: practice mode and compete mode. Results from runs played in compete mode are counted towards a player’s position on a leaderboard, which is visible to other players, while results from runs in practice mode are visible only to the player. Paired sample t-tests showed that players (*N* = 2303) using both practice and compete modes on the same day performed statistically significantly worse while in practice mode compared to compete mode (hits per second: *t* = 71.533, *df* = 6364, *p*-value < 1e-6; hit rate: *t* = 23.721, *df* = 6364, *p*-value < 1e-6). However, we assume that Gridshot runs performed in practice mode could contribute to learning. Therefore, for each player, *K* runs per day and *K* runs total (between 7/1/20-1/30/21) were based on Gridshot runs where mode was either practice or compete, while performance metrics (hit rate and hits per second) were limited to runs played in compete mode.

Our main dependent variable for performance accuracy was hit rate, proportion of shots that hit targets, and our main dependent variable for motor acuity was hits per second, the product of speed (attempted shots per second) and performance accuracy (hit rate). Speed and accuracy were computed separately for each 60 second run of Gridshot (Figure 2, individual data points), along with score (Figure 2, colors). Each quartile of score exhibits a speed - accuracy tradeoff; speed is higher when accuracy is lower and vice versa. Runs with higher scores have correspondingly both higher speed and higher accuracy. Motor acuity increases as the product of speed and accuracy. This is represented in Figure 2 along the 45-degree diagonal, up and to the right.

**Figure 2.**
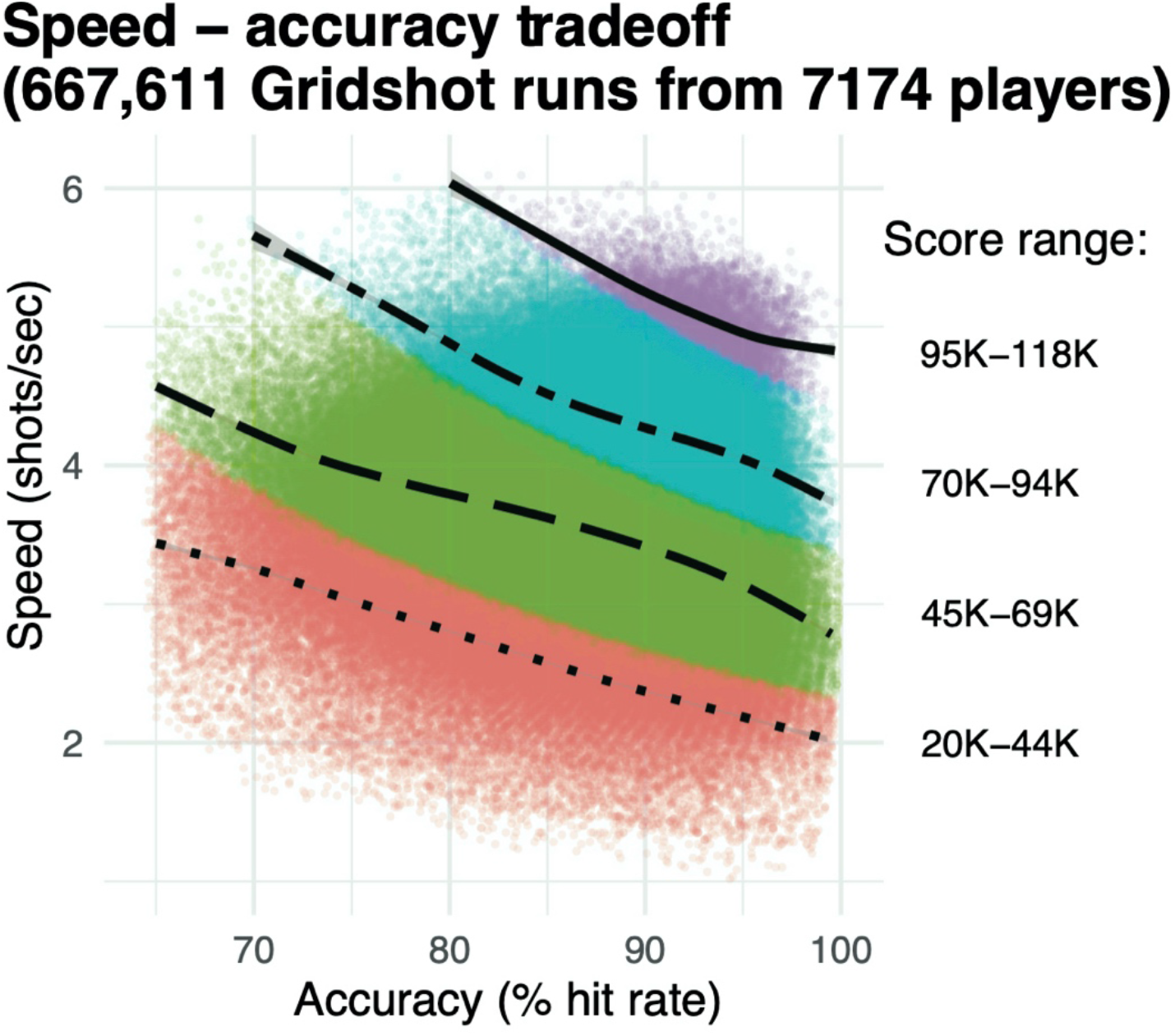
Speed - accuracy tradeoff. Individual data points, speed (shots per second) and accuracy (hit rate) for each individual 60 second run of Gridshot. Colors, score quartiles. Curves, locally weighted regression fits per score quartile.

#### Does motor performance accuracy and motor acuity improve across days of gameplay?

Specifically, we asked how hit rate, our measure of performance accuracy, and hits per second, our measure of motor acuity, improved across the number of days of gameplay.

#### How does the amount of practice affect motor performance accuracy, motor acuity, and retention?

Specifically, we asked how the amount of learning on day *M*, defined as the number of runs on a given game day, impacted the player’s improvement in performance from baseline (i.e., hits per second on run 1 of day 1).

The number of players decreased with each subsequent run and each subsequent day, because some players stopped playing. Consequently, to examine patterns of learning within a day and retention of learning to the following day, the data were filtered to exclude performance data from days for which the previous calendar date had been skipped and where *N* players < 200 for each data point. The remaining data set included 5144 players’ Gridshot data. For ease of exposition, we limited our analysis of learning within a day to runs 1-25 on days 1-5. Since different players played different numbers of runs on each day, the sample sizes for each data point were uneven (Day 1, *N* = 5144; Day 5, *N* = 221).

We also asked how the amount of learning on day *m*-1 impacted retention on day *m*. Retention was calculated as 100(*S*_*m*,1_−*S*_*m*−1,1_)/(*S*_*m*−1,25_ –*S*_*m*−1,1_), where *S*_*m,x*_ is the increase in score from baseline for run *x* on day *m*. Thus, retention, as operationally defined here, represents the magnitude of learning (performance improvement) that is retained from the end of one day to the start of the next, normalized by the previous day’s total magnitude of learning. We restricted the analysis to consecutive days of game play within days 2 through 21, and furthermore to consecutive days with *N* ≥ 20 players. We removed outlier values (prob > 0.95) of the resulting retention metric, for a given day *m*, based on median absolute deviation using the outliers package in R (Komsta, 2011). We then fit the relationship between retention and day number with a nonparametric regression model that fits a polynomial function, based on local fitting (locally weighted scatter plot smoother a.k.a. loess regression) (Cleveland, 1979; Komsta, 2011), using the loess function from the stats package in R (degree = 2, span = 0.75) and plotted the results (R Core Team, 2021).

#### How much should gamers practice on a given day to maximize improvements in motor acuity?

Specifically, we characterized how improvements in hits per second from run *k*, day *m* to run *k*, day *m*+1 for *k* = 1, 2, and 3, varied as a function of the number of runs on day *m*. After calculating improvement per player from one day to the next, we removed outlier values (prob > 0.95) of improvement, for a given value of *k*, based on median absolute deviation, (Komsta, 2011). For each value of *k* (1 - 3), we fit the relationship between number of runs the previous day and median change in hits per second using the loess function from the stats package in R (degree = 2, span = 0.75) and plotted the predicted values based on the model. We restricted our analyses to runs up to 100 on days 1-21 (first 3 weeks) of game play to maintain a large player sample size per data point. The remaining data set included Gridshot data from 3461 players.

#### Exploring which factors of gameplay contribute to improvements in motor acuity

To characterize the phenomena observed with the methods described above and to understand the possible contributions of various factors to motor learning over time, within the context of this first-person shooter task, we fit repeated measures linear mixed models (growth model), separately for hits per second and hit rate as outcome variables. The models accounted for repeated (correlated) measures within players as well as different performance levels and practice behaviors between players.

Data were limited to Gridshot runs with day number ≤60 to maintain a large sample size for each day number in the test set. Based on pilot analyses indicating that improvements are non-linear, we transformed day number and median runs per day into log(day number) and log(median runs per day). Fixed effects included start values for each performance metric on day 1, log(median runs per day), days from last play, log(day number), and interactions between all fixed effects. We allowed for random intercepts and slopes for each player. Due to the large sample size of the training data set (*N* players = 5712), tests for normality of outcome measures were not necessary.

We built and validated a pair of models (one for hit rate and one for hits per second) via backward stepwise elimination followed by cross-validation. Data were split into train and test sets (80/20) using the groupdata2 package in R (Olsen, 2021), evenly distributing players based on total days played. We started with model variants that included maximal combinations of regressors. These models were fit to the training data (*N* players = 5712) using the R package lme4 (algorithm “REML”, optimiser “nloptwrap”, link function “identity”, family “gaussian”) (Bates et al., 2015; Kuznetsova et al., 2017). The R package lmerTest was used to perform backward stepwise elimination from each maximal model to reach a final model based on likelihood ratio tests. The final pair of models were validated using the test set (*N* players = 1427).

To evaluate predictor variables for multicollinearity, prior to model testing, correlation coefficients among all potential predictor variables were calculated. All pairwise correlation coefficients were -0.07 to +0.07. Marginal and conditional R^2^ were similar for train and test sets for each of the two models developed to explain factors that contribute to variance in motor learning within the context of Gridshot, indicating that the models were not over-fitted. (Table 1)

**Table 1:** Model Validation.

| Model | Conditional R<sup>2</sup> | Marginal R<sup>2</sup> | RMSE |
| --- | --- | --- | --- |
| HPS train | 0.934 | 0.708 | 0.126 |
| HPS test | 0.935 | 0.721 | 0.126 |
| Hit Rate train | 0.797 | 0.256 | 0.224 |
| Hit Rate test | 0.806 | 0.249 | 0.217 |

The resulting model structures are reported in Table 2, which contains coefficient estimates (“Estimates”) as a measure of effect size and a *p*-value via Satterthwaite’s degrees of freedom method. Tables include marginal and conditional R^2^ for mixed models (Nakagawa & Schielzeth, 2013), and variance (among players) computed in the R package sjPlot (Lüdecke, 2021).

**Table 2:** Model Performance.

|  | median day hps |  |  | median day hit rate |  |  |
| --- | --- | --- | --- | --- | --- | --- |
| *Predictors* | *Estimates* | *CI* | *p* | *Estimates* | *CI* | *p* |
| (Intercept) | 0.74 | 0.71 – 0.78 | <0.001 | 1.35 | 1.29 – 1.40 | <0.001 |
| start_hps | 0.79 | 0.77 – 0.81 | <0.001 |  |  |  |
| log(day_number) | 0.24 | 0.23 – 0.26 | <0.001 | 0.34 | 0.31 – 0.36 | <0.001 |
| log(median_runs) | 0.1 | 0.07 – 0.14 | <0.001 | 0.04 | 0.02 – 0.06 | <0.001 |
| days_fromlastplay | 0 | -0.01 – 0.00 | 0.131 | -0.05 | -0.06 – -0.04 | <0.001 |
| start_hps * log(day_number) | -0.06 | -0.07 – -0.05 | <0.001 |  |  |  |
| start_hps * log(median_runs) | -0.03 | -0.05 – -0.00 | 0.018 |  |  |  |
| log(day_number) * log(median_runs) | 0.04 | 0.03 – 0.06 | <0.001 | 0.01 | 0.01 – 0.02 | <0.001 |
| start_hps * days_fromlastplay | 0 | -0.01 – -0.00 | 0.003 |  |  |  |
| log(day_number) * days_fromlastplay | -0.01 | -0.02 – -0.01 | <0.001 | -0.03 | -0.04 – -0.03 | <0.001 |
| log(median_runs) * days_fromlastplay | -0.01 | -0.02 – -0.00 | 0.001 | 0 | 0.00 – 0.00 | 0.043 |
| (start_hps * log(day_number)) *<br>log(median_runs) | -0.01 | -0.02 – -<br>0.00 | **0.005** |  |  |  |
| (start_hps * log(day_number)) *<br>days_fromlastplay | 0 | 0.00 –<br>0.00 | **0.044** |  |  |  |
| (start_hps * log(median_runs)) *<br>days_fromlastplay | 0.01 | 0.00 –<br>0.01 | **0.007** |  |  |  |
| (log(day_number) *<br>log(median_runs)) *<br>days_fromlastplay | -0.01 | -0.01 – -<br>0.00 | **0.001** |  |  |  |
| (start_hps * log(day_number) *<br>log(median_runs)) *<br>days_fromlastplay | 0 | 0.00 –<br>0.01 | **0.007** |  |  |  |
| start_hit_rate |  |  |  | 0.39 | 0.36<br>- 0.42 | **<0.001** |
| start_hit_rate *<br>log(day_number) |  |  |  | -0.15 | -0.16 - -<br>0.14 | **<0.001** |
| start_hit_rate *<br>days_fromlastplay |  |  |  | 0.01 | 0.01<br>- 0.02 | **<0.001** |
| (start_hit_rate *<br>log(day_number)) *<br>days_fromlastplay |  |  |  | 0.01 | 0.01<br>- 0.01 | **<0.001** |
| Random Effects |  |  |  |  |  |  |
| $\sigma^2$ | 0.02 | | | 0.06 | | |
| $\tau_{00}$ | 0.06 player ID | | | 0.15 player ID | | |
| $\tau_{11}$ | 0.01 player<br>ID.log(day_number) | | | 0.02 player<br>ID.log(day_number) | | |
| $\rho_{01}$ | 0.74 player ID | | | 0.65 player ID | | |
| ICC | 0.77 |  |  | 0.73 |  |  |
| N | 5712 player ID |  |  | 5712 player ID |  |  |
| Observations | 42635 |  |  | 42635 |  |  |
| Marginal R<sup>2</sup> / Conditional R<sup>2</sup> | 0.708 / 0.934 |  |  | 0.256 / 0.797 |  |  |

## Results

### Does motor performance accuracy and motor acuity improve across days of gameplay?

We used hit rate, percentage of shot attempts that successfully hit a target, to measure how motor performance accuracy improved over 100 days of gameplay. A greater hit rate indicated that the player was more accurate in hitting the target. Players’ (*N* = 7174) hit rate ranged between 65.0% and 99.5% on day 1, exhibiting a wide distribution among players. This heterogeneity in performance was likely attributed to the wide range of gaming experience among players on the Aim Lab platform, which ranged from amateur to professional.

We found a significant increase in hit rate over time (Figure 3A). Specifically, median hit rate on day one was 86.4% and on day 100 was 89.6% (*t* = -4.17, *df* = 104.28, *p*-value = 3.11e-05). Improvements in hit rates were also non-linear (R^2^ non-linear: 0.020, R^2^ linear: 0.017), perhaps because many players started to reach a ceiling level of performance. The average rate of increase was 0.032% per day, exhibiting a very modest rate of improvement in motor performance accuracy over time.

**Figure 3.**
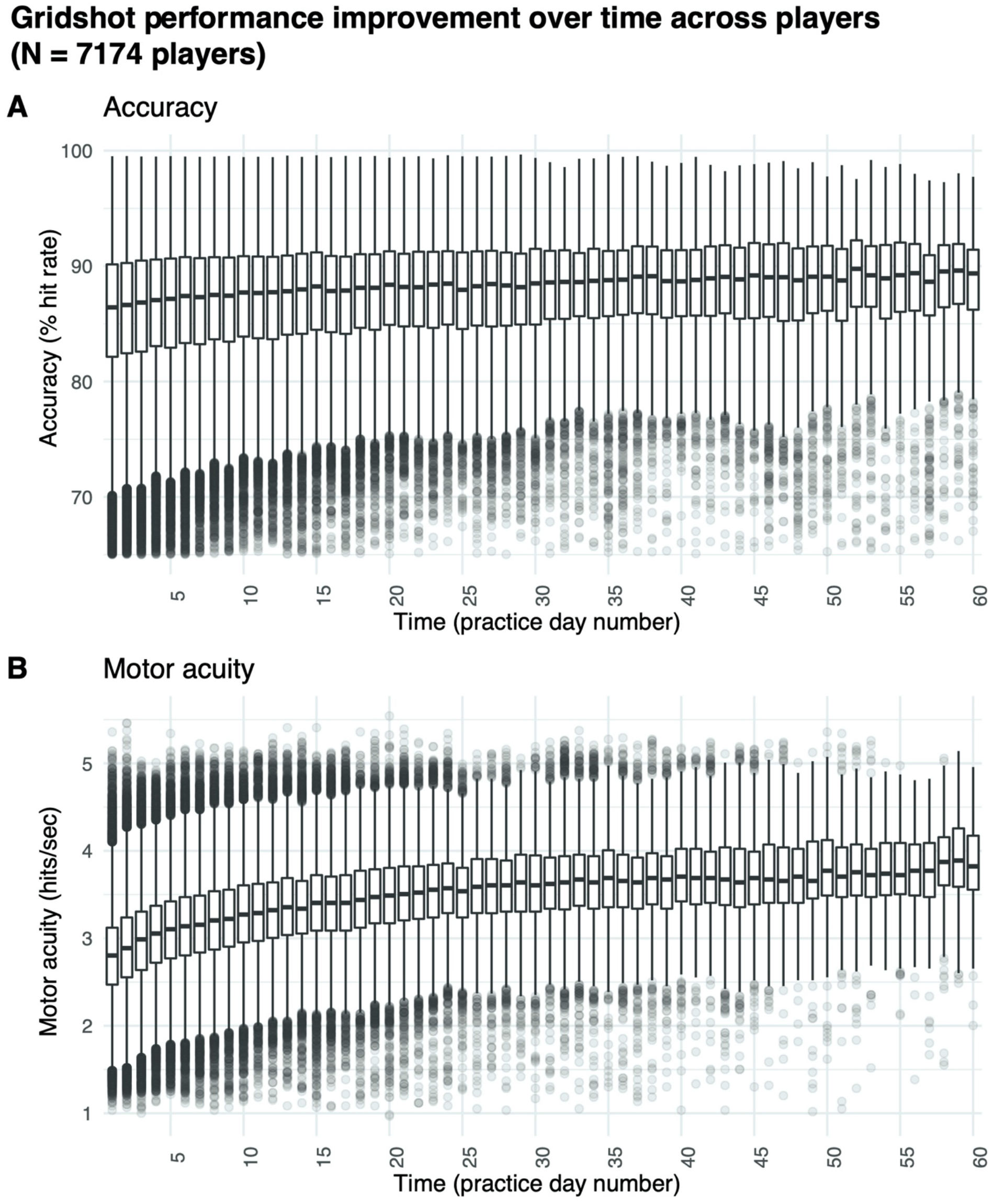
Improvements in motor skill over 100 days of shooting practice. **(A)** Changes in motor performance accuracy (hit rate: percentage of shots attempted that hit targets) and **(B)** motor acuity (hits per second: number of accurate shots that hit targets every second). Box plots denote the median (thick horizontal lines), quartiles (1st and 3rd, the edges of the boxes), and extrema (min and max, vertical thin lines).

Improvements in hit rate, however, may not be the most sensitive measure of motor learning for two reasons: first, measures of performance accuracy may readily hit a ceiling or a floor – a concern that is also present in our own dataset. Second, because changes in performance only occur early in learning, they may measure improvements in task-irrelevant dimensions, such as familiarization with which button to click or even posture in the gaming chair, rather than improvements in the motor skill of interest (e.g., shooting a target).

Measures of motor acuity, like hits per second (number of targets successfully destroyed per second), offer a more sensitive measure of motor learning. These measures typically exhibit a greater range of improvement both early and late in learning, and as such, can better capture improvements in motor execution rather than only improvements in task-irrelevant dimensions. Hits per second ranged between 1.04 and 5.36 on day 1, also exhibiting large heterogeneity amongst players. To put this range in perspective, ten of the recent top Gridshot performers in Aim Lab had a median hits per second of 6.25, indicating that the median performance among players in our sample is far from ceiling.

Most critically, we found large improvements in motor acuity over time (Figure 3B). Specifically, the median hits per second was 2.80 on day 1, whereas it was 4.00 on day 100 (*t* = -28.577, *df* = 104.4, *p*-value < 1e-6). As for hit rate, improvements in hits per second were also non-linear (R^2^ non-linear: 0.208, R^2^ linear: 0.255), showing a trend similar to in-lab studies studying motor skill acquisition (Telgen et al., 2014; Yang et al., n.d.) and validating our online approach.

Together, we observed improvements in both motor performance accuracy and motor acuity during first-person video gameplay, providing a proof-of-concept in studying motor learning outside the lab.

### How does the amount of practice affect motor performance accuracy, motor acuity, and retention?

We first asked how practice influences how much gamers improve in their motor performance accuracy (i.e., hit rate) from their baseline performance on their first run on day one. We limited our analyses to players who completed five consecutive days of shooting practice with at least 25 runs on each day (each data point in Figure 4 represents more than 200 players).

**Figure 4.**
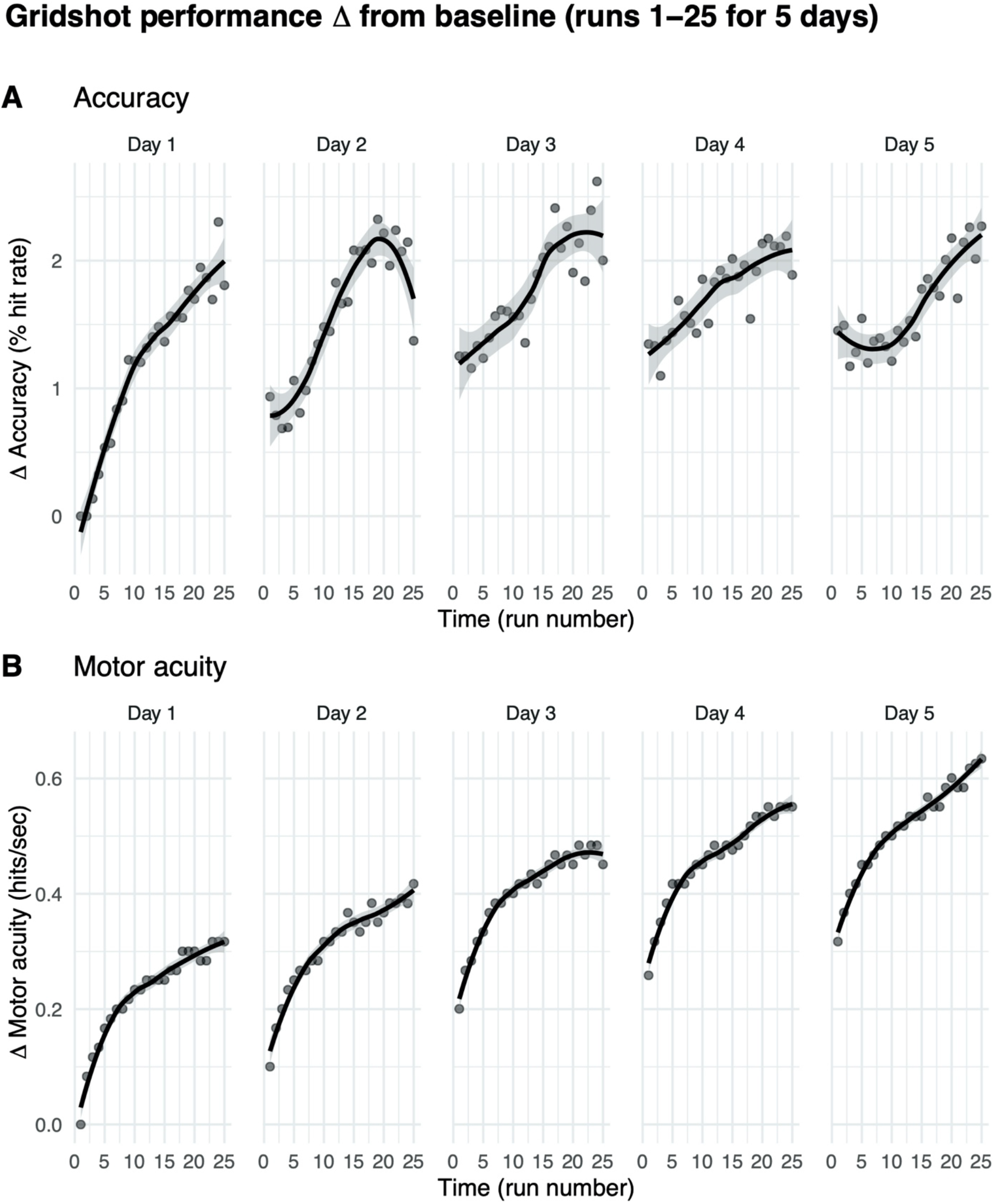
Motor performance exhibits within-day and across-day improvements. **(A)** Motor performance accuracy (hit rate). **(B)** Motor acuity (hits per second). Curves, locally weighted regression fitted to the data. Shaded error bars denote the 95% confidence interval.

Hit rates increased from baseline within each game day (Figure 4B). On day 1, for instance, within-day performance change from baseline per run increased significantly from 0% on run 1 (baseline, by definition) to 2.091% on run 25 (*t* = -8.713, *df* = 638, *p*-value < 1e-6). The rate of increase on day 1 was moderate, exhibiting a 0.087% improvement in hit rate per run. These data suggest that more practice runs within the same gameday increase shooting accuracy.

We then asked whether performance improved across these five days. Average improvements in hit rate increased across days 1 to 3 (day 1: 0.23 +/- 2.12, day 2: 1.25 +/- 6.16, day 3: 1.6 +/- 6.58.), but there was no evidence for any additional increase from days 3 to 5 (*t* = -0.39866, *df* = 3761.7, *p*-value = 0.345). That is, average hit rate increase on each day plateaued, exhibiting an asymptotic mean improvement in hit rate of 1.7% +/- 6.37%. While these data suggest that gamers were becoming more accurate shooters with greater within and across-day practice, motor performance accuracy seemed to saturate, hitting an upper bound with as little as five days of practice. These data also highlight how hit rate cannot capture the richer aspects of motor learning that occur over a prolonged period of practice (e.g., 100 days, see Figure 3B).

Changes in motor acuity (i.e., measured by hits per second) provided a better, more sensitive index of motor skill acquisition both within each day of practice and across days of practice (Figure 4B). On day 1, change from baseline in hits per second increased significantly from 0 (baseline, by definition) to 0.323 (*t* = -27.134, *df* = 638, *p*-value < 1e-6). That is, players were able to hit one more target roughly every three seconds. These within-day improvements increased with more practice runs in a non-linear manner (marginal R^2^ linear (0.120) < marginal R^2^ nonlinear (0.324)). The rate of increase on day 1 was quite fast, exhibiting a 0.013 hits per second improvement per run. Indeed, 25 runs of practice helped gamers hit targets more accurately and more rapidly.

There was a rapid improvement in hits per second over the first few runs on each successive day (Figure 4B). We hypothesize that this “warm-up” effect is, in part, due to motor adaptation (see Discussion).

Across days, change from median hits per second on the first run to the 25^th^ run on each day increased from 0.32 +/- 0.3 on day 1 to 0.63 +/- 0.35 on day 5. In contrast to hit rate, motor acuity improvement did not show signs of plateauing; instead, hits per second continued to improve over each day.

We then asked how much of the improvements in motor performance accuracy and motor acuity are retained across days? As shown in Figure 5A and B, the first run on day *m*+1 was greater than the first run on day *m*, suggesting that some of the performance/acuity gains were retained (see *Methods* for details).

**Figure 5.**
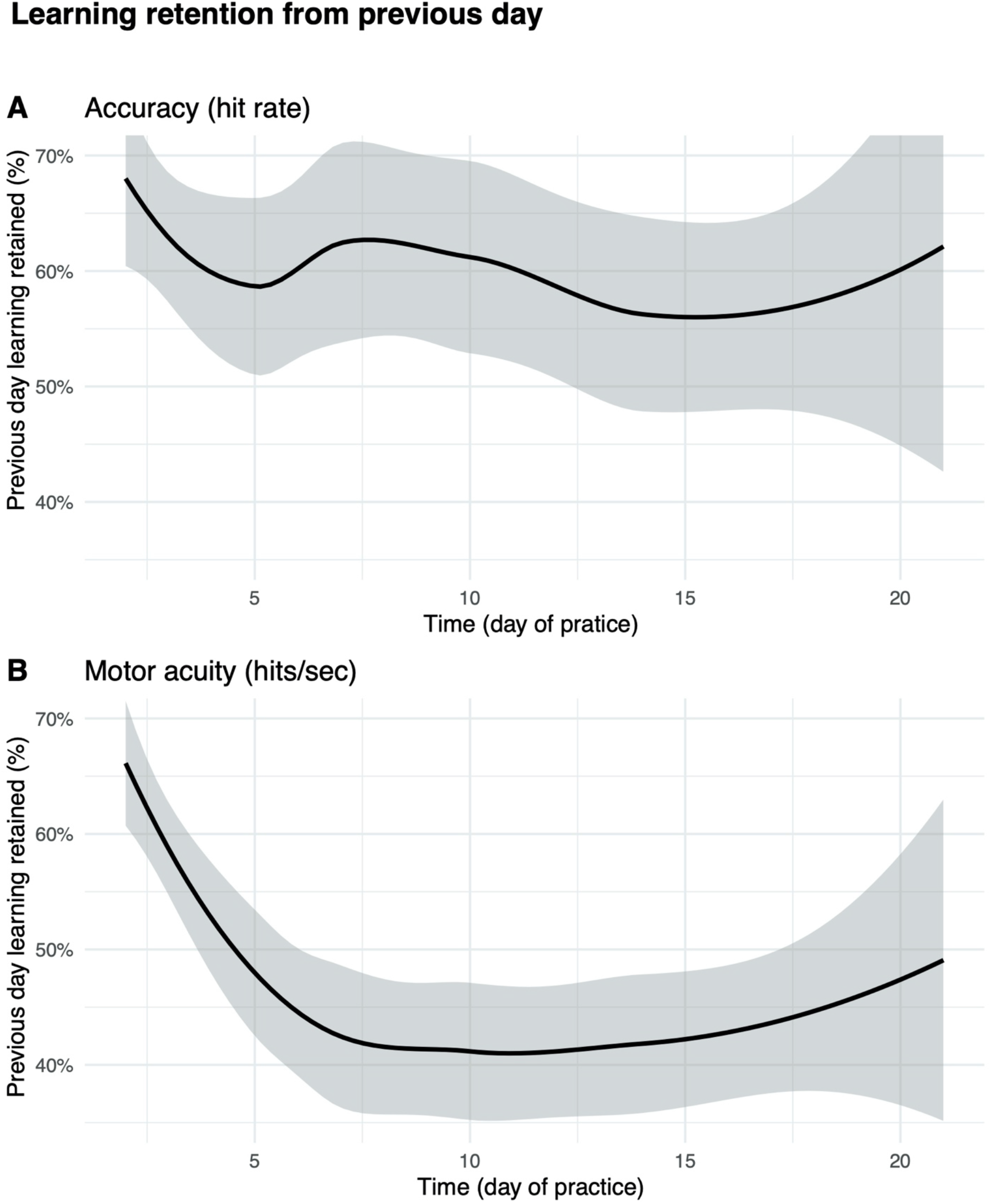
Percent improvement on day m retained on day m + 1. **(A)** Motor performance accuracy (hit rate) **(B)** Motor acuity (hits per second). Curves, locally weighted regression fitted to the data. Shaded error bars denote the 95% confidence interval.

Performance accuracy (hit rate) retention was greatest in the early days of practice (Figure 5A). For instance, participants were able to retain as much as 67% +/- 79% improvements in hit rate from day 1 to day 2. The percent value of retention did not decrease drastically over subsequent days. This is likely attributable to the small absolute within-day gains in performance accuracy, as players reached a ceiling level of performance accuracy (hit rate) as early as day 3 (Figure 4A).

We next examined how motor acuity (hits per second) was retained from one day to the next (Figure 5B). On day 2, players retained 65% +/- 52% of what they learned from day 1. Percent retention declined across days, plateauing at 40% after day 10. This effect not only implies that motor acuity was still exhibiting within-day improvements (also supported by Figure 4B), but also that improvements in motor acuity could be carried over to the next day.

### How much should gamers practice on a given day to maximize improvements in motor acuity?

While more practice on a given day may yield greater improvements in hits per second, does more practice always mean that these improvements are retained? Here, we reasoned that there could be a sweet spot in motor learning. Whereas too little practice would be detrimental to retention since little was learned in the first place, too much practice may also be detrimental due to mental/physical fatigue (Branscheidt et al., 2019). To explore this question, we calculated how much motor acuity improved across days *m* to *m*+1 as a measure of how long gamers practiced on day m.

We found that the benefit of additional practice exhibited a non-monotonic function (Figure 6). The greatest improvements in motor acuity were evident with 75 runs when comparing the first run on two consecutive days, and 90% learning benefit was achieved by practicing 49 runs, which roughly corresponds to 50 minutes of gameplay. A similar pattern held when comparing run 2 and run 3 performance on two consecutive days, showing maximum benefit of learning with 60 (90% benefit after 41 runs) and 57 runs of practice (90% benefit after 33 runs), respectively. Taken together, this non-monotonic function provides strong evidence for diminishing returns on practice, similar to those of previous studies (Stafford & Dewar, 2014). As noted above, we observed a rapid improvement in motor acuity over the first few runs on each successive day (Figure 4B), a “warm-up” effect that we hypothesize is due to motor adaptation (see Discussion).

**Figure 6.**
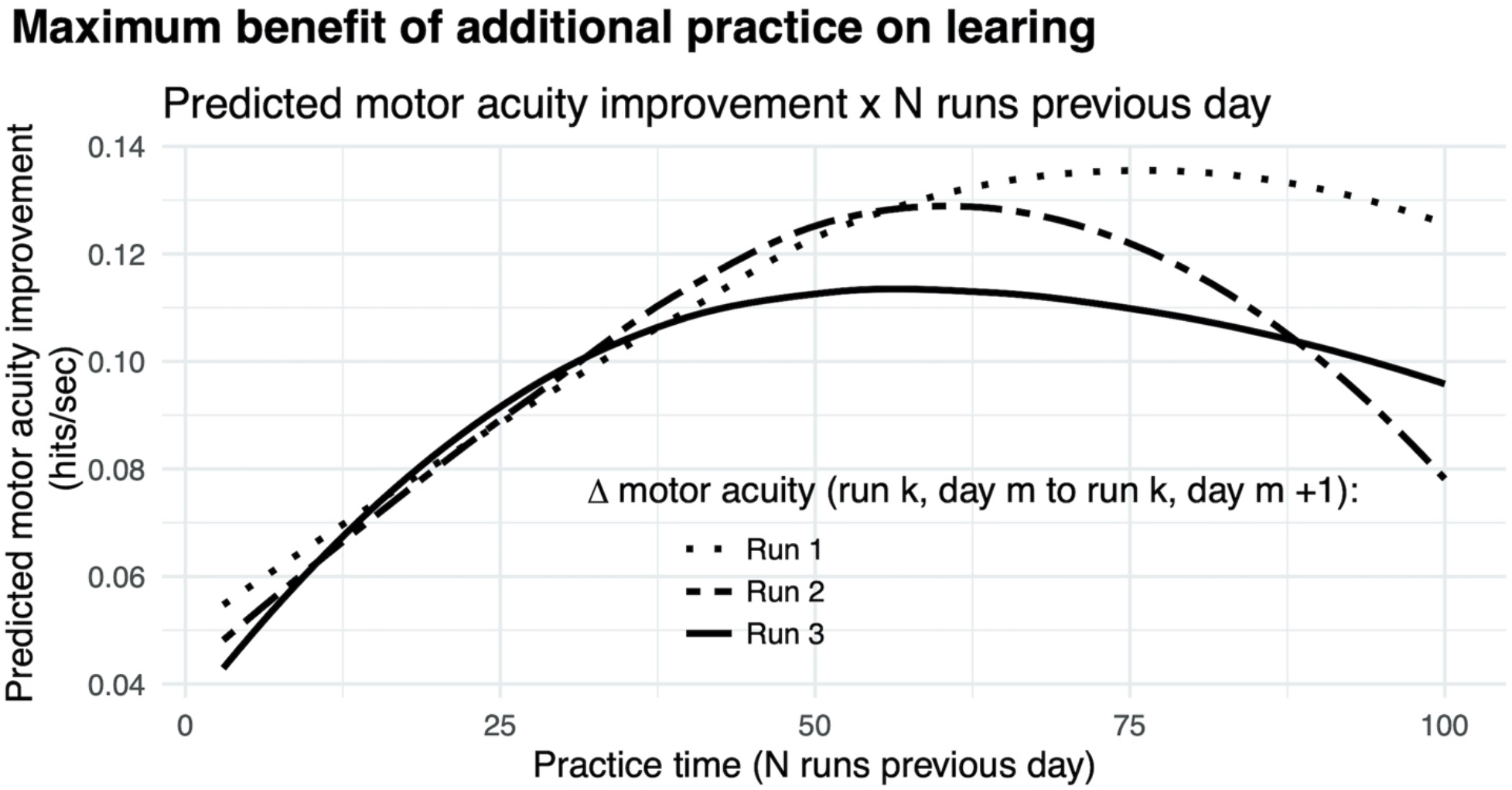
Diminishing returns on motor acuity with more practice on the previous day. Curves, model predictions for improvements in motor acuity (hits per second) on run 3 (solid), run 2 (dashed), and run 1 (dotted) as a function of number of runs in the previous day.

After 3 runs of warm-up on every day, the maximal improvement in motor acuity across days is seen with about an hour of practice (Figure 6, dashed curve). With no warm-up, considerably more practice (75 minutes) on any given day is needed to achieve maximal benefit the following day (Figure 6, solid curve). We infer that the additional practice is needed to compensate for the lack of warm-up.

### Exploring which factors of gameplay contribute to gamers’ improvements in motor acuity?

We took an exploratory approach to the data by building the most parsimonious models (via stepwise regression and cross-validation), using all known, measurable factors of gameplay to explain their contributions to gamers’ improvements in motor performance accuracy (hit rate) and motor acuity (hit rate per second) over time.

The best-fitting model that converged and best explained change in performance accuracy (hit rate) included main effects for baseline performance accuracy (i.e., median hit rate on day 1), number of days of practice (log(day number)), amount of practice within a day (log(median runs per day)), and number of days skipped between practice days, with interactions effects between baseline performance accuracy x log(day number), baseline performance accuracy x days from last play, log(day number) x days from last play, log(day number x log(median runs per day), baseline performance accuracy x log(day number) x days from last play. Because each player started playing with a different level of performance accuracy, we allowed each player to have their own slope and intercept (Table 2).

The significant negative interaction effect of day number on baseline motor performance accuracy indicates the better a player’s starting hit rate, the less their hit rate increased per additional day of practice; or, the magnitude of the positive effect of a unit increase in start hit rate on predicted learning was decreased by each successive day number. While skipped days (days from last play) had statistically significant interaction effects with other variables, along with that of the main effect for skipped days, the effect sizes (coefficients) were quite small, meaning skipping days between practice sessions has a small negative effect on motor performance accuracy that could likely be overcome with additional practice.

To provide some intuition behind our model, we illustrate how our model predicts motor performance accuracy for a typical player. A typical player with modest accuracy (a baseline hit rate of 85.67% on day 1), who plays 10 min of Gridshot recreationally per day (10 runs, without skipping any days of practice), may be predicted to have a hit rate of 86.50% on day 10, 86.73% on day 20, 86.85% on day 30, and 86.94% on day 40. If instead they played 20 runs of Gridshot per day, the same player would have predicted values of 86.82% hit rate on day 10, 87.13% on day 20, 87.31% on day 30, and 87.43% on day 40. The predicted improvements in motor performance accuracy are quite modest.

The model that best explained change in motor acuity (hits per second) included main effects for baseline motor acuity (i.e., median hit rate on day 1), number of days of practice (log(day number)), amount of practice within a day (log(median runs per day)), number of days skipped between practice days, and all potential interaction effects, with independent slope and intercept for each player (Table 2).

As was the case for hit rate, there was a significant negative interaction effect of day number on start value for hits per second, meaning the greater a player’s starting value for hits per second, the less increase there was in motor acuity per additional day of practice. Similarly, there was a negative coefficient for the interaction effect between median runs per day and start value for hits per second, implying that additional practice within a day had less of an effect on the magnitude of learning for players who were more skilled to begin with. Again, skipped days (days from last play) had statistically significant interaction effects with other variables in the explanatory model, as well as a statistically significant main effect, but with negligible coefficients.

For example, a typical player with a start value of 2.78 hits per second on day 1 who plays 10 runs of Gridshot per day, skipping no days of practice, would have predicted values of 3.17 hits per second on day 10, 3.29 on day 20, 3.35 on day 30, and 3.39 on day 40. If instead they played 20 runs of Gridshot per day, the same player would have predicted values of 3.22 hits per second on day 10, 3.35 on day 20, 3.42 on day 30, and 3.47 on day 40.

Model predictions are summarized in Figure 7A & B. Rate of learning is a function of performance level at the start of learning (hits per second and hit rate start value), median runs of Gridshot per day, and to a lesser extent, number of days between practice sessions. The plots demonstrate the non-linear relationship between additional runs of practice per day and increase in learning per day as well as the non-linear pattern of learning over time, regardless of number of runs per day. In addition, the model of hit rate (Figure 7A) demonstrates that motor performance accuracy is predicted to decrease if it is greater than 88% (could be a “sweet spot” of learning: see discussion). The difference between marginal and conditional R^2^ for both models (hits per second 0.708 / 0.934; hit rate 0.256 / 0.797) estimates the extent to which unmeasured variation among players contributes to variance in each model. For example, the low marginal R^2^ (0.256) and higher conditional R^2^ (0.797) for the hit rate model indicates that most of the variance in hit rate is explained by differences among players, while most of the variance in hits per second can be explained by the known fixed effects.

**Figure 7:**
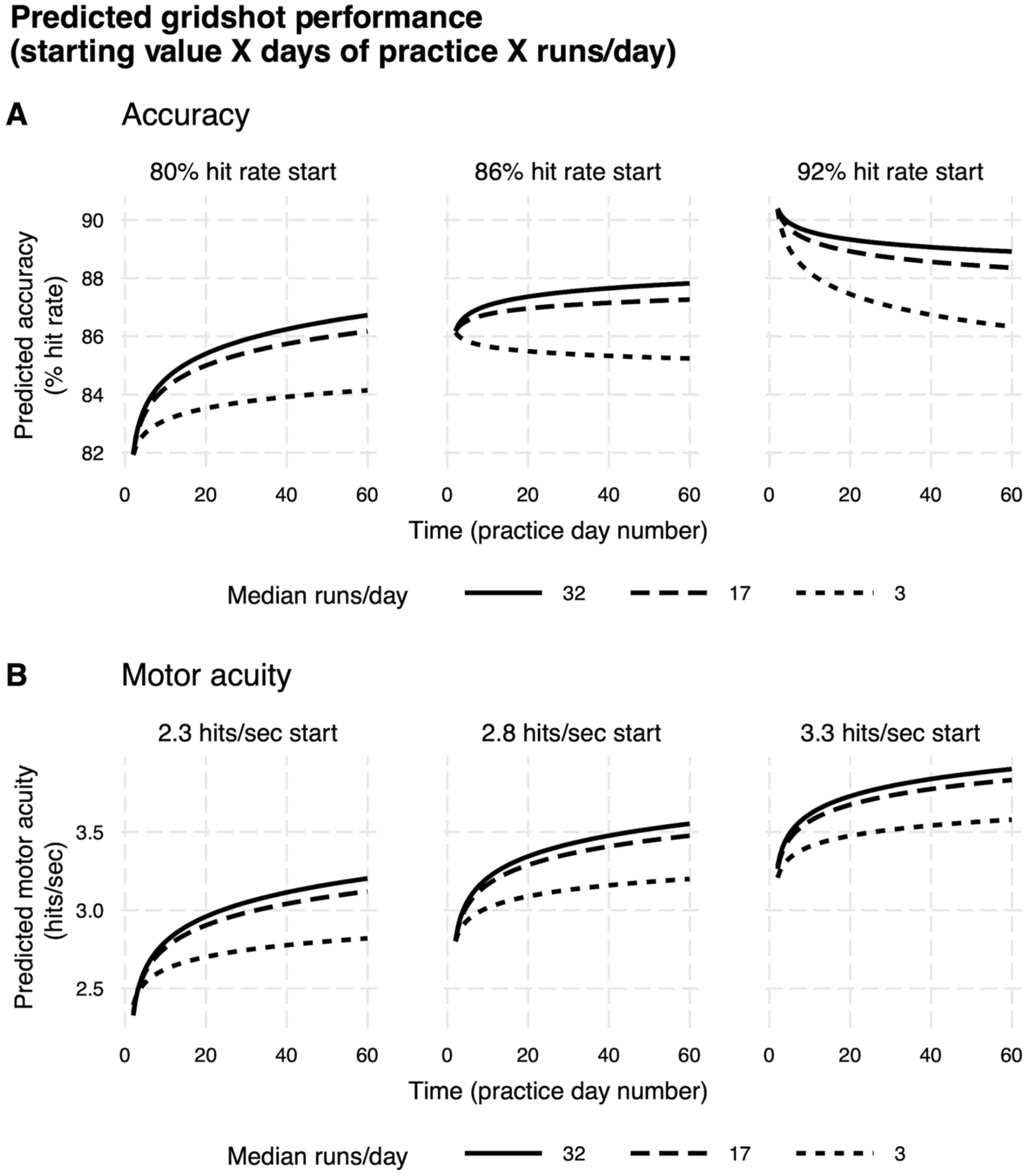
Predicted learning curves. **(A)** Motor performance accuracy (hit rate). **(B)** Motor acuity (hits per second). Different panels delineate model predictions based on differences in baseline motor performance. Curves denote differences in the amount of practice on each day.

A more accurate way to model outcome measures on which there is a ceiling effect is an asymptotic regression as time goes to infinity (Chambers & Noble, 1961; Stevens, 1951). Our implementations of asymptotic regression models indicated that our data collection time period did not extend long enough for player performance to reach an asymptote for either outcome measure. However, our regression models provide operational estimates of the contributions of our predictor variables to outcome measures within the confines of the study time period.

## Discussion

While motor learning enables us to successfully interact with our world, it is seldom studied outside the laboratory. As a result, research has focused on how either simple or somewhat contrived motor skills can be quickly acquired in a well-controlled environment (e.g., circle drawing or compensating for mirrored visual feedback; (Hadjiosif et al., 2020; Kasuga et al., 2015; Shmuelof et al., 2012; Telgen et al., 2014)). Moreover, the population of study is typically homogenous and recruited via financial incentives (Henrich et al., 2010). As such, whether these observations from the lab generalize to the acquisition of much more complex real-world motor skills remains an open question. Here, using a first-person shooter game platform (Aim Lab), we provide a proof-of-concept of studying motor skill acquisition in a real-world setting. We not only amassed a large sample of motivated gamers who span a wide range of expertise, but also observed how their motor skills progressed longitudinally over 100 days. We found that while their shooting accuracy saturated within a few days, motor acuity continued to show improvements (i.e., shifts in the speed-accuracy tradeoff). Furthermore, these improvements were modulated by the duration of practice, with diminishing returns after roughly 30 - 60 minutes of practice.

This proof-of-concept study thus showcases a powerful new approach to study motor learning in the wild, demonstrating in the context of motor learning research, the scientific value of data collected on a large scale in non-controlled conditions. Limitations of this study included factors either not measurable or not taken into account: Aim Lab tasks the player might have played, other than Gridshot; time of day tasks were played; number of sessions contributing to a single day of player data; time spent playing other video games; player equipment and internet speed differences; player demographic, lifestyle, and health behaviors such as sleep, caffeine intake, handedness or age.

Although motor learning is comprised of a diversity of learning processes, a large proportion of studies in the field have focused specifically on motor adaptation in response to visual or somatosensory perturbations, in which the goal of the participant is to return to baseline levels of performance through recalibration of sensorimotor mappings (Krakauer et al., 2019). We observed a rapid improvement in motor acuity (hits per second) over the first few runs on each successive day (Figure 4B) that we hypothesized is due to motor adaptation. Specifically, Aim Lab (like most games of this genre) enables players to set the mouse sensitivity; mouse sensitivity has units of °/cm and determines the amount of rotation in the virtual environment of the game for each centimeter of mouse movement. The mouse sensitivity when playing the game is typically incommensurate with mouse usage in other applications (e.g., navigating the desktop to start the game, and navigating the Aim Lab user interface to initiate Gridshot). Hence the need for motor adaptation to recalibrate movements with respect to the mouse sensitivity setting.While adaptation is key to successful movement in the face of changing states and environments,the goal of movement practice outside the lab, for example, in sports or rehabilitation, is oftentimes to far exceed baseline levels of performance. Such improvements in motor acuity are accomplished not through motor adaptation but rather through motor skill acquisition, the gradual improvement in our ability to select and execute appropriate actions.

Let’s now consider how our results relate to the three distinct stages of motor learning proposed by Fitts and Posner (Fitts & Posner, 1979) (see Introduction). Motor performance accuracy, as measured by hit rate (i.e., number of targets hit/total shooting attempts), is likely most sensitive to improvements during the cognitive stage. Once the movement requirements are understood – for instance, which keypress to make or how one should sit comfortably in their chair – hit rate may quickly saturate. Indeed, this phenomenon is present in our data, with hit rates reaching a ceiling around five days, similar to the findings from previous lab-based motor learning studies (McDougle & Taylor, 2019; Shmuelof et al., 2012). For instance, in visuomotor learning studies where the participant is required to hit the target with their perturbed cursor (e.g., 30° clockwise rotation), performance often reaches an upper bound within a few reaches (Albert et al., 2021; Harris, 1963; Hutter & Taylor, 2018; Kornheiser, 1976; Langsdorf et al., 2021; McDougle & Taylor, 2019; Morehead et al., 2015; Stratton, 1896; J. A. Taylor et al., 2014). In other words, participants are quick to figure out how to nullify the perturbation by reaching in the opposite direction. As a result, after these initial trials only marginal improvements in motor accuracy are observed within a single session of practice (Bond & Taylor, 2015; Huberdeau et al., 2019; Neville & Cressman, 2018). The difference in time to saturation between GridShot and visuomotor rotation (days vs minutes) likely reflects the increased complexity and task demands of our gaming task.

In contrast to motor performance accuracy, *motor acuity*, as measured by the number of targets being hit per second, may be more sensitive to improvements during the association stage of motor learning. As evident in our data, hits per second saturates more slowly, and continues to improve for some players over as long as 100 days. Despite performing at a ceiling level in accuracy, players still make considerable improvements in how fast they are moving. Returning to the visuomotor learning literature, a similar analogy can also be made to studies performed in the lab. For example, following introduction of a visuomotor rotation, participants can learn to accurately hit the target with the perturbed cursor, but at the cost of much slower reaction times as compared to baseline reaches (Benson et al., 2011; Fernandez-Ruiz et al., 2011; McDougle & Taylor, 2019; Wilterson & Taylor, 2021). Therefore, even after several days of practice, there remains ample room for improving motor acuity by reducing reaction times (Wilterson & Taylor, 2021). Taken together, unlike cruder measures of motor performance (e.g., accuracy), motor acuity, which takes into account both speed and accuracy, may serve as a more sensitive measure to quantify the progression of motor learning over long periods of practice (Shmuelof et al., 2012, 2014).

This contrast between motor performance accuracy and motor acuity has also been noted by several others (Krakauer et al., 2019). For instance, in a seminal study by Shmuelof et al, participants were instructed to draw circles as accurately and quickly as they could within a tight boundary (Shmuelof et al., 2012, 2014). Improvements in motor skills were quantified by shifts in the speed-accuracy tradeoff function describing the number of circles drawn within a given amount of time. They found that while participants could perform the task relatively accurately within one day of practice, improvements in the speed-accuracy tradeoff and coordination (e.g., smoothness of the motor trajectories) required over five days of practice. Similar to our study, their measure of motor acuity best captured improvements in motor learning over time (also see: (Telgen et al., 2014)). Thus, it appears that mastering a motor skill requires prolonged practice over long timescales, whether in the context of learning arbitrary visuomotor mappings (Hardwick et al., 2019), rolling cigars (Crossman, 1959), or, as we show here, gaming.

We were also interested in the critical ingredients that influence the rate of motor skill acquisition. From our exploratory analysis, we found that baseline performance, number of days of practice (day number), median number of runs per day, and the consistency of practice schedule (quantified by number of intervening days between practice sessions) impact improvement in motor acuity and motor performance accuracy (Table 2). Notably, we found that better performance at baseline predicts less improvement with each additional day of practice (Figure 7). Similarly, proficient players benefitted less with additional practice within a given day. Additionally, we observed diminishing returns on motor learning with increased practice (Newell & Rosenbloom, 1981; Rubin-Rabson, 1941; E. D. Ryan, 1965) (Figure 6). Specifically, while roughly one hour of gameplay allows for maximal retention across days, practicing more than this did not improve learning, possibly due to fatigue (Branscheidt et al., 2019). Taken together, our analyses provide several important constraints on motor skill acquisition during video game play, which motivate future experiments to isolate and characterize each factor in detail.

In addition to improvements in motor performance accuracy and motor acuity, we also saw other principles of motor learning in the wild. Specifically, we observed that players retained ∼65% (on average) of their learning from day 1 to day 2. These values are larger than those of previous motor adaptation studies conducted in the lab, with participants retaining ∼40% of their learning over consecutive days of practice (Joiner & Smith, 2008; Telgen et al., 2014). Yet, 65% retention is smaller than retention values following learning a skill de novo (i.e., mirror reversal) (Telgen et al., 2014; Wilterson & Taylor, 2021). As such, we speculate that our intermediate retention value reflects the fact that multiple learning systems are engaged during Gridshot, including recalibration of a learned skill as well as acquiring a new skill.

While there were similarities between the overall approach and findings observed in the present study as compared to many lab-based studies, there were also some fundamental differences that are likely attributable to the differences between tasks. First, tasks used in the lab often involve simple, predictable movements (e.g., center-out reaching, serial reaction time sequence learning tasks, drawing circles). Here, our first-person shooting game, in contrast, requires a complex set of unpredictable movements (e.g., shooting three unknown targets presented sequentially as fast and as accurately as possible). That is, successful gamers not only need to click the correct mouse button and make the appropriate keypresses to shoot, but they also need to immediately plan and execute a set of discrete movements towards unexpected target locations. Arguably, most movements made in daily life are more akin to the longer planning horizons required in first-person shooter games, demanding that multiple actions need to be learned and simultaneously retrieved in short term memory (Gallivan et al., 2018). It is likely these qualitative differences in task demands that contributed to the steady improvements in motor acuity across multiple days of Gridshot practice, rather than quickly hitting a performance ceiling as in most motor learning tasks studied in the lab.

Second, whereas in-lab tasks typically focus on isolating and dissociating one learning mechanism from another (Avraham et al., 2021; Hegele & Heuer, 2010; Kim et al., 2018; Leow et al., 2018; Morehead et al., 2017; Nikooyan & Ahmed, 2015; Pellizzer & Georgopoulos, 1993; Tsay et al., 2020; Tsay, Haith, et al., 2021; Tsay, Kim, et al., 2021), our task lies on the opposite end of the spectrum, where a wide range of learning processes are likely involved. For instance, cognitive strategies related to how to play the game (e.g., planning sequences of movements), motor adaptation (recalibrating movements with respect to mouse sensitivity), and skill learning (i.e., increasing the speed of both wrist and finger motions without sacrificing accuracy) all contribute to success in the game. Moreover, gamers are most certainly also learning via reward and punishment to determine which movement strategies to retain and which ones to abandon. That is, there is likely an interaction between reinforcement learning and working memory (Collins, 2017; Collins et al., 2014; Collins & Cockburn, 2020)). While the analytical tools required to extract how each learning process contributes to motor learning still need to be fully fleshed out, gaming nonetheless provides an ideal way to gather rich data for such multi-dimensional analyses.

Perceptual learning may also contribute to gains in Gridshot performance due to gamers being able to identify and locate relevant targets quickly and accurately. Perceptual learning is optimized through prolonged training, often with significant and long-lasting performance benefits (for reviews, (Censor et al., 2012; Dosher & Lu, 2017; Maniglia & Seitz, 2018; Sagi, 2011; Sasaki et al., 2010; Seitz & Dinse, 2007; Watanabe & Sasaki, 2015). The perceptual demands for Gridshot are modest compared to most perceptual tasks (detecting and localizing targets), but the pace at which the task is performed (hitting up to 6 targets per second) surely stresses the capability of the visual system.

We speculate that ∼85% hit rate optimizes both learning and task engagement. We observed that players exhibit a steady level of performance accuracy (∼85%) throughout learning, even as motor acuity continues to increase due to faster speed (shots per second). One could imagine a different outcome in which performance accuracy increases to nearly 100% without increasing speed. Participants in a previous study likewise voluntarily selected an intermediate difficulty level while learning (Baranes et al., 2014). Errors in learning are beneficial for human learning (Metcalfe, 2017), and 85% accuracy is nearly optimal during the training of machine learning algorithms as well (Wilson et al., 2019). To maintain 85% hit rate while learning, players may adopt a strategy in which they increase their speed slightly, causing accuracy to decrease, and then practice at that new speed until their performance accuracy recovers before increasing their speed again.

The future of studying motor learning through video games is promising (Anguera et al., 2013; Chen et al., 2018; Stafford & Dewar, 2014; Stafford & Vaci, 2021; Tsay, Ivry, et al., 2021). Gaming provides a unique medium to investigate how motor skills are developed through intrinsic motivation, rather than financial incentive. For instance, previous studies examining motor learning during neurorehabilitation have suggested that intrinsic motivation is a critical ingredient for motor retention (Kleim & Jones, 2008; Tsay & Winstein, 2020; Winstein et al., 2014; Winstein & Varghese, 2018). The statistically significant within-player performance difference between Gridshot runs played in practice (scores visible only to the player) vs compete (scores visible to other players) modes in this study demonstrates the effect that motivation can have on player performance in a motor learning task. In a similar vein, gaming naturally lends itself to longitudinal studies, rather than being limited to cross-sectional studies (Crossman, 1959), affording greater within-subject controls and insights into the individual differences of motor expertise (Anderson et al., 2021). Gaming also provides a convenient way to amass a wealth of moment-to-moment kinematic data. Recent studies have shown how much kinematics can reveal, ranging from variables related to decision making (Freeman & Ambady, 2010; Song & Nakayama, 2008, 2009) to those related to one’s reward expectations (Sedaghat-Nejad et al., 2019; Summerside et al., 2018). Future iterations of Aim Lab task-based studies will focus on collecting and analyzing these fine-grained kinematic data, which will provide deeper insights into how complex motor skills can be acquired and refined in an ecological setting, outside the traditional laboratory.

## Author Contributions

DH assisted with study design and manuscript preparation. WM designed and implemented the study apparatus and tasks. JL carried out data analyses and assisted with study design and manuscript preparation. HK and JT assisted with manuscript preparation. All authors read and approved the final manuscript.

## Acknowledgments

The authors would like to thank the Aim Lab users whose playing time and performance data have contributed to this work.

## Data Availability Statement

The raw data for this manuscript were acquired for commercial purposes. The data are not scheduled to be made publicly available because they are proprietary.

